# Deconvolution of single-cell multi-omics layers reveals regulatory heterogeneity

**DOI:** 10.1101/316208

**Authors:** Longqi Liu, Chuanyu Liu, Andrés Quintero, Liang Wu, Yue Yuan, Mingyue Wang, Mengnan Cheng, Lizhi Leng, Liqin Xu, Guoyi Dong, Rui Li, Yang Liu, Xiaoyu Wei, Jiangshan Xu, Xiaowei Chen, Haorong Lu, Dongsheng Chen, Quanlei Wang, Qing Zhou, Xinxin Lin, Guibo Li, Shiping Liu, Qi Wang, Hongru Wang, J. Lynn Fink, Zhengliang Gao, Xin Liu, Yong Hou, Shida Zhu, Huanming Yang, Yunming Ye, Ge Lin, Fang Chen, Carl Herrmann, Roland Eils, Zhouchun Shang, Xun Xu

## Abstract

Integrative analysis of multi-omics layers at single cell level is critical for accurate dissection of cell-to-cell variation within certain cell populations. Here we report scCAT-seq, a technique for simultaneously assaying chromatin accessibility and the transcriptome within the same single cell. We show that the combined single cell signatures enable accurate construction of regulatory relationships between *cis*-regulatory elements and the target genes at single-cell resolution, providing a new dimension of features that helps direct discovery of regulatory patterns specific to distinct cell identities. Moreover, we generated the first single cell integrated maps of chromatin accessibility and transcriptome in human pre-implantation embryos and demonstrated the robustness of scCAT-seq in the precise dissection of master transcription factors in cells of distinct states during embryo development. The ability to obtain these two layers of omics data will help provide more accurate definitions of “single cell state” and enable the deconvolution of regulatory heterogeneity from complex cell populations.

---

The rapid proliferation of single cell sequencing technologies has greatly improved our understanding of heterogeneity in terms of genetic, epigenetic and transcriptional regulation within cell populations^1^. We, and others, have developed single-cell whole genome^2^, exome^3, 4^, methylome^5^ and transcriptome^6, 7^ technologies and applied these approaches to analyzing the complexity of cell populations in tumorigenesis, developmental process and cellular reprogramming^8^. Meanwhile, single-cell epigenome techniques, including single cell ChIP-seq^9^, ATAC-seq^10, 11^, DNase-seq^12^ and Hi-C^13, 14^, have been developed to decipher histone modifications, transcription factor (TF) accessibility landscapes, and 3D chromatin contacts, respectively, in single cells. These techniques provide important information on regulatory heterogeneity by assessing chromatin structure across various cell types.

Measuring the epigenomic and transcriptomic characteristics of single cells is important for understanding the maintenance and conversion of cell fates, as well as manipulating cell fates into different lineages^15^. The regulation of these processes involves sequential events including the binding of TFs to *cis-*regulatory elements (CREs) and the recruitment of chromatin regulators, resulting in changes of chromatin structure and activation or repression of cell type specific genes^15^. Single-cell ATAC-seq and RNA-seq represent a great opportunity to study how TFs and epigenomic features induce transcriptional outcomes that influence cell fate determinations. For example, combined analyses of datasets by these two approaches have enabled characterization of subtypes in mouse tissues^16^ or during human hematopoietic differentiation^17^. However, it still remains challenging to integrate the two approaches experimentally in individual cells, thus hampering a full understanding of regulatory association between these two layers. Here, we present scCAT-seq (**s**ingle-**c**ell **c**hromain **a**ccessibility and **t**ranscriptome **seq**uencing), a technique that integrates single-cell ATAC-seq and RNA-seq to measure chromatin accessibility (CA) and gene expression (GE) simultaneously in single cells. scCAT-seq employs a mild lysis approach and a physical dissociation strategy to separate the nucleus and cytoplasm of each single cell. Thereafter, the supernatant cytoplasm component is subjected to the Smart-seq2 method as described previously^7^. The precipitated nucleus is then subjected to a Tn5 transposase-based and carrier DNA-mediated protocol to amplify the fragments within accessible regions (Fig. 1a and **Supplementary Methods**). Beyond parallel CA and GE profiling in the same single cell, scCAT-seq will be particularly useful for analyzing samples when the amount of input material is limited.

**Figure 1.**
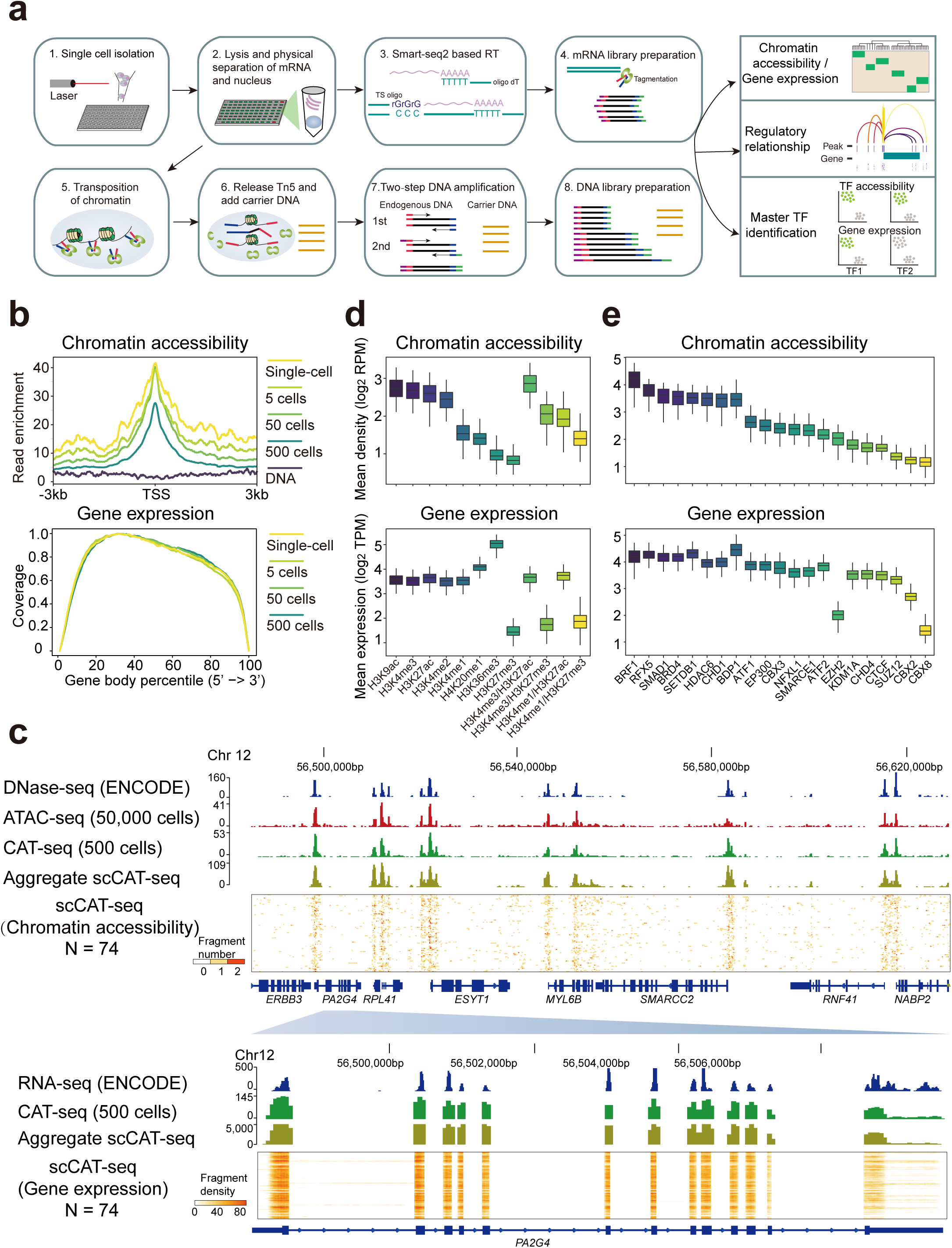
scCAT-seq provides an accurate genome-wide measure of both chromatin accessibility and gene expression. (**a**) Overview of the scCAT-seq protocol. (**b**) Top panel: chromatin accessibility read enrichment around the transcription start site (TSS). Bottom panel: coverage of mRNA reads along the body of transcripts. Titration series (one single-cell, 5 cells, 50 cells, 500 cells) were marked by the indicated colours. All profiles were generated using the scCAT-seq protocol with the indicated number of cells as input. (**c**) A representative region showing a consistent pattern of chromatin accessibility and gene expression across datasets generated using different number of input cells. The bulk ATAC-seq track was generated using 50,000 K562 cells. The DNase-seq and bulk RNA-seq data of K562 cells were downloaded from ENCODE. The scCAT-seq tracks are chromatin accessibility (upper) and gene expression read density (bottom) from a total of 74 K562 single cells. (**d**) Top panel: mean chromatin accessibility read density around regions that are enriched by the indicated individual or combined histone modifications. Bottom panel: mean expression level of genes associated with regions that are enriched by the indicated individual or combined histone modifications. (**e**) Top panel: mean chromatin accessibility read density within regions that are bound by the indicated transcription factors. Bottom panel: mean expression level of genes associated with regions that are bound by the indicated transcription factors.

## Results

### Simultaneous profiling of accessible chromatin and gene expression in single cells

We applied scCAT-seq to the K562 chronic myelogenous leukemia cell line, which has been widely used in the ENCODE project. We sorted single cell and multi-cell samples (e.g., 500 cells) into wells of 96-well plates using flow cytometry. Empty wells were used as negative control. Samples were then processed using the scCAT-seq protocol. qPCR analysis confirmed the successful capture of single cell nuclei during library preparation (Supplementary Fig. 1a). We generated combined CA and GE profiles from a total of 192 samples. Of the 176 single cell profiles, 74 (42.0%) of them passed both CA and GE data quality control criteria (Supplementary Fig. 1b and **Supplementary Methods**).

For scCAT-seq-generated CA data, we obtained an average of 2.1 x 10^5^ uniquely mapped, usable fragments from single cells (**Supplementary Table 1** and Supplementary Fig. 1c,d). Similar to bulk ATAC-seq^18^, the CA fragments show fragment-size periodicity corresponding to integer multiples of nucleosomes (Supplementary Fig. 1e) and are strongly enriched on accessible regions (Fig. 1b and **Supplementary Table 1**). We found that about 9% of the fragments were mapped to the mitochondrial genome (Supplementary Fig. 1f) which is largely reduced in comparison to standard bulk ATAC-seq studies (typically over 30%)^18^. Pearson correlation analyses revealed our single-cell profiles could reproduce features of bulk profiles (Supplementary Fig. 1g). In comparison to the published scATAC-seq profiles by Buenrostro *et al.*^10^, we obtained a higher number of usable fragments per single cell but with lower signal-to-noise ratio (Supplementary Fig. 1h). However, the correlation between single cells increases remarkably (Supplementary Fig. 1h), suggesting that scCAT-seq is able to capture the chromatin features more accurately.

For mRNA data generated by scCAT-seq, we obtained an average of 4.6 million reads covering over 8000 genes (GENCODE v19, TPM > 1), which is comparable to published scRNA-seq profiles by Pollen *et al*.^19^ (Supplementary Fig. 1j and **Supplementary Table 1**). Consistent with published Smart-seq profiles, our mRNA data showed full coverage of the transcript body (Fig. 1b), enabling identification of transcript isoforms and not merely gene expression quantification. The aggregate profile was close to the RNA-seq profile obtained from 500 cells (Pearson correlation value > 0.9, Supplementary Fig. 1i), suggesting that scCAT-seq is able to accurately quantify GE of single cells. The density of CA and GE reads of all single cells surrounding a constitutively accessible region showed that scCAT-seq data recapitulate major features obtained by separately performed bulk ATAC-seq and RNA-seq (Fig. 1c).

GE regulation is associated with the structure of the CREs (e.g., histone modifications, DNA methylation) and the binding of *trans*-factors (e.g., TFs, epigenetic modifiers)^20^. Therefore, we examined the overall distribution of single-cell CA fragments across different genomic contexts, as well as the expression levels of the putative regulated genes. We observed that the CA fragments were enriched at CREs with active histone modifications (e.g., H3K27ac, H3K9ac and H3K4me3), whereas repressive or inaccessible regions (e.g., H3K27me3 and H3K36me3-associated regions) showed lower fragment density (Fig. 1d). We also observed other association patterns between CA and GE. For example, we found low levels of CA fragments on H3K36me3-associated regions but high levels of GE. This is not surprising because H3K36me3 is known to be enriched on the active gene body which is occupied by nucleosomes and rendered inaccessible^20^. Notably, genes with bivalent marks (coenrichment of H3K4me3 or H3K4me1 and H3K27me3) showed similar level of accessibility as active genes (co-enrichment of H3K4me3 or H3K4me1 and H3K27ac, but lack of H3K27me3), and both of them showed higher levels of accessibility than inactive genes (enrichment of H3K27me3, but not H3K27ac, H3K4me1 and H3K4me3). Conversely, the expression levels of bivalent genes were remarkably lower than active genes and were similar to those of inactive genes. We also investigated the distribution of CA fragments across genomic contexts bound by different TFs and found an overall consistent pattern between CA and GE level. Notably, we observed substantial decrease of expression levels of genes associated with binding of EZH2 while the accessibility level showed just a moderate change (Fig. 1e). This pattern is similar to that of bivalent genes and is consistent with the role of EZH2 which, as part of the repressive polycomb complex, catalyzes H3K27me3. Thus, the combined signatures from scCAT-seq well reflect known processes well and are useful to assess the transcriptional state of genes within different genomic contexts. This approach is undoubtedly of high value for many biological applications, for example, studying the heterogeneous transition of bivalent genes during development or cellular reprogramming.

We further validated our approach by generating different batches of scCAT-seq profiles from two additional ENCODE cell lines: HeLa-S3 cervix adenocarcinoma and HCT116 colorectal carcinoma cell lines **(Supplementary Table 1)**. To test the feasibility of scCAT-seq in real tissue samples, we also generated profiles from two lung cancer patient-derived xenograft (PDX) models **(Supplementary Table 1)**. One is derived from a moderately differentiated squamous cell carcinoma patient (PDX1) and the other one from a large-cell lung carcinoma patient (PDX2). Principal components analysis (PCA) on both CA and GE profiles resulted in separation of cells from different origin (Supplementary Fig. 2a,b). A comparison of our datasets with published profiles revealed that the differences across protocols and batches had a substantially smaller effect than difference across cell types (Supplementary Fig. 2c,d).

### Establishment of regulatory relationships between CREs and genes in single cells

Next, we explored the dynamic associations between the two omics layers across single cells. We first tested the correlation between accessibility level of single CREs and their expression of the putative target genes in each of the three cell lines, and the hypothetical cell population merged from them. As expected, we identified remarkably more positive correlations (Pearson correlation > 0; FDR < 10%) than negative correlations (Supplementary Fig. 3a), which is consistent with the known relationship between CA and GE in bulk profiles^21^.

An earlier study showed the co-variability of accessibility between CREs across single cells defines regulatory domains highly concordant with observed chromosome compartments, which provides an alternative approach to the discovery of regulatory links^10^. However, it still remains impossible to directly infer the transcriptional outcomes of each chromatin accessible region. Given the overall positive correlation between CA and GE, we reasoned that the co-variability between accessibility of individual elements and expression of genes could enhance discovery of regulatory links that influence transcription. To this end, while employing the reported strategy using scATAC-seq^10^ (strategy 1, Fig. 2a), we proposed two additional strategies for inferring regulatory relationships (strategy 2 and 3, Fig. 2a). For strategy 1 and 2, regulatory relationships between chromatin accessible regions and target genes were identified based on scATAC-seq and scCAT-seq data, respectively. Based on scATAC-seq data, regulatory relationships for every gene were assigned when the Spearman correlation of the accessibility of CREs located at the promoter and distal peaks was above 0.25 (strategy 1, Fig. 2a and **Supplementary Methods**). Likewise, for the scCAT-seq data, the regulatory links were assigned if the Spearman correlation between the GE and the accessibility of distal CREs was above 0.25 (strategy 2, Fig. 2a and **Supplementary Methods**. However, these regulatory relationships are defined across all cells. In order to more accurately depict the regulatory relationship between chromatin and genes, in strategy 3, single-cell-specific regulatory relationships between genes and their nearby accessible regions were assigned using the scCAT-seq data as follows: i) identification of active TFs for every cell by SCENIC^22^ using the normalized GE matrix; ii) identification of active accessible regions by matching the binding motifs of active TFs to accessible chromatin regions; and iii) assignment of regulatory relationships after applying a Wilcoxon test to determine if the presence of a nearby active accessible region was associated with a significant change in the target GE (p-value < 0.05) (Fig. 2a and **Supplementary Methods**.

**Figure 2.**
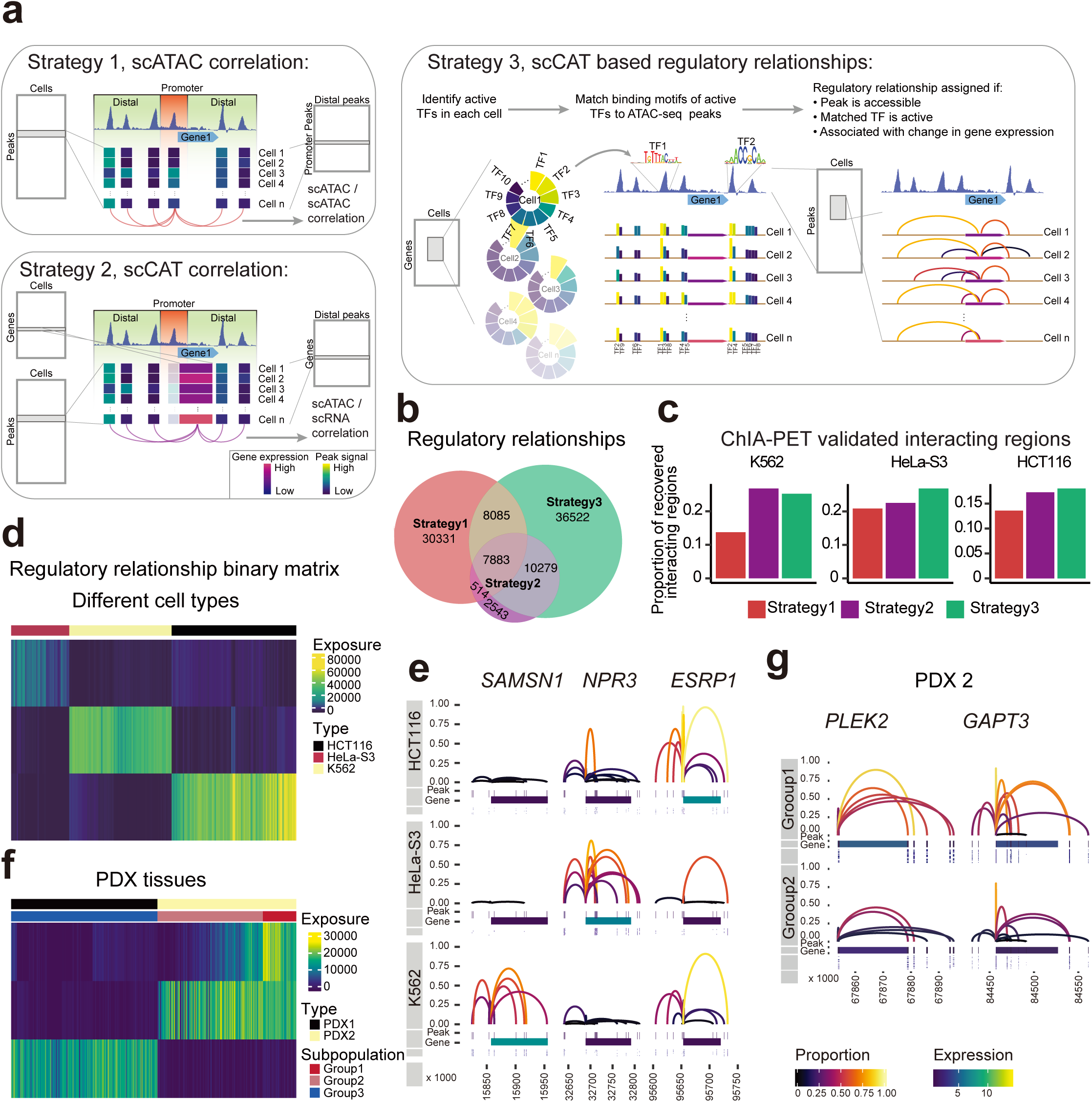
Inferring regulatory relationships between CREs and genes by scCAT-seq. (**a**) Overview of three strategies for inferring regulatory relationships. Strategy 1: regulatory links for every gene were assigned when the Spearman correlation of the signal of peaks located at the promoter and distal peaks was above 0.25. Strategy 2: the regulatory links were assigned if the Spearman correlation between the gene expression and the signal of distal peaks was above 0.25. Strategy 3: active transcription factors for every cell were identified by SCENIC, then active regions were identified by matching the binding motifs of active transcription factors to accessible regions. Then regulatory relationships were assigned after applying a Wilcoxon test to determine if the presence of a nearby active accessible region was associated with a significant change in the target gene expression (p-value < 0.05). (**b**) Venn plot showing the number of overlapping regulatory relationships identified by the three strategies. (**c**) Proportion of ChIA-PET validated regulatory relationships identified by the three strategies in K562 (left), HeLa-S3 (middle) and HCT116 (right) single cells. (**d** and **f**) Heatmaps showing exposure scores of all cells to each signature identified by the NMF clustering of regulatory relationship binary matrix in cell lines (**d**) and PDX **(f)**. The exposure score represents the contributions of the signatures to the different samples. **(e** and **g)** Regulatory relationships for the indicated genes in single cell groups of the cell lines **(e)** and PDX2 **(g)**. Each panel contains three tracks: the top track shows the regulatory relationship between one peak and the gene (linking them with an arch), where the height and colour of the arch show the proportion of cells that share the regulatory relationships; the middle track shows the genomic location of the gene and the associated peaks, where the colour of the gene shows the mean expression in each cell type; the bottom track shows the accessible states (on and off) for each peak in each single cell.

By applying the 3 strategies to single cells of the 3 cell lines, we found that strategy 3 identified the largest number of regulatory relationships (62,769), compared to strategy 1 (46,813) and strategy 2 (21,219) (Fig. 2b). Over 1/3 of the regulatory relationships from scATAC-seq based method (strategy 1) were shared by those from scCAT-seq based method (strategy 2 and 3), suggesting strong synergistic effects between regulation at chromatin and transcriptome levels. Nevertheless, although a similar correlation approach was used in strategies 1 and 2, strategy 2 identified a lower number of regulatory relationships, suggesting a possible decoupling between accessibility at the promoter and the expression of the gene. Notably, we also observed a large fraction of regulatory relationships specifically identified by each method, which suggests that different information can be obtained from single-omics and combined analysis.

To assess the accuracy of the regulatory links inferred by each method, we next counted the regulatory relationships that could be verified by chromatin interaction analysis by paired-end tag sequencing (ChIA-PET)^23^. Encouragingly, using the ChIA-PET interactions of the three widely used cell types (K562, HeLa-S3 and HCT116)^24^, we observed higher proportion of validations in scCAT-seq based method (strategy 2 and 3) than that in scATAC-seq based method (strategy 1) in all 3 cell types (Fig. 2c). These suggest that the co-variability between CA and GE layers could better reflect higher-order chromatin structure than co-variability between CREs. One explanation is that regulatory relationships inferred from scATAC-seq may result from either chromatin interactions or from co-binding of master TFs without interaction, while those inferred from scCAT-seq could be considered to be “functional” regulatory relationships as include information from both chromatin interactions and co-binding of master TFs. Therefore, based on the largest number of validated regulatory relationships, strategy 3 outperformed the other strategies (hereafter, the “regulatory relationship” indicates those identified only by strategy 3). The distribution of distance between each pair of peak and gene in all regulatory relationships showed higher enrichment in proximal regions than distal regions (Supplementary Fig. 3b), suggesting that GE tends to be regulated by proximal elements which is consistent with earlier findings^25^.

To assess whether the regulatory relationships in each single cell reflect cell type-specific features, we generated a binary matrix where columns represent single cells, and rows represent all identified regulatory relationships between accessible sites and genes, and the entries indicate the on or off state of each regulatory relationship in each cell. We applied a non-negative matrix factorization (NMF) method, implemented in the R package Bratwurst^26^, to decompose the matrix into different signatures that could distinguish single cell identities.As expected, NMF clustering of the regulatory relationships identified signatures containing numerous cell type-specific regulatory relationships, resulting in clear separation of the 3 cell types (Fig. 2d,e and Supplementary Fig. 3c). For example, *SAMSN1* is a known oncogene, preferentially expressed in the blood cancer, multiple myeloma^27^. We observed highly specific regulatory relationships around *SAMSN1* in K562, a myelogenous leukemia cell line (Fig. 2e), revealing a strong association between its expression and accessibility of CREs. This observation again reconfirmed the importance of epigenetic mechanisms during progression of tumors. Likewise, we generated regulatory relationship matrix for single cells from PDX tissues and clustering of the matrix clearly separated these two type of cells (Fig. 2f,g and Supplementary Fig. 3d). Interestingly, we also observed a subpopulation of cells showing specific regulatory relationships in PDX2 (Fig. 2f,g), likely reflecting the regulatory heterogeneity present in real tissues.

### Integrated single-cell epigenome and transcriptome maps of human pre-implantation embryos

We next explored the potential of scCAT-seq in the characterization of single cell identities in continuous developmental processes. The human pre-implantation embryo development is a fascinating time that involves dramatic changes in both chromatin state and transcriptional activity. However, it has only been investigated at either the chromatin or the RNA level due to the lack of truly integrative approaches^28^. By using clinically discarded human embryos (**Supplementary methods**), we generated scCAT-seq profiles for a total of 110 individual cells, and successfully obtained 29 quality-filtered profiles from morula stage and 43 from blastocyst stage (success rate 65.5%) (Fig. 3a, Supplementary Fig. 4a and **Supplementary Table 1**). To explore the regulation relevant to each stage, we identified ∼100K regulatory relationships and generated a matrix of regulatory relationships across all single cells as described above. NMF clustering analysis of the matrix showed separation of all single cells into two main groups (group 1 and 2), corresponding to these two stages (Fig. 3b). The heatmap of exposure scores to each signature revealed activation of regulatory relationships of pluripotency markers (such as NANOG and KLF17) in morula, and trophectoderm (TE) markers (such as CDX2 and GATA3) in blastocyst stage^28^ (Fig. 3b,c and Supplementary Fig. 4b,c), which strongly suggests that the expression of these markers is activated/maintained by epigenomic states^28^.

**Figure 3.**
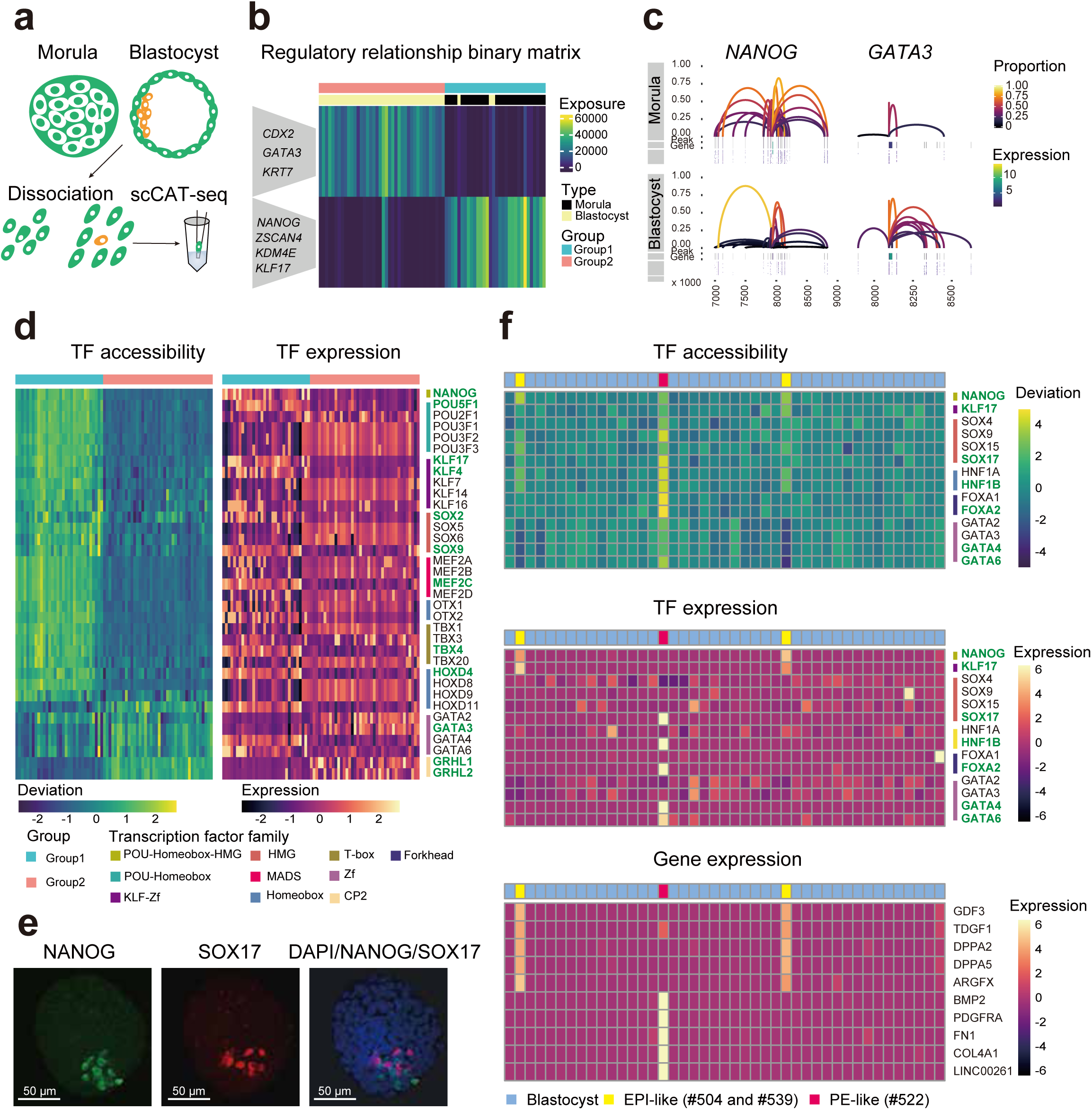
scCAT-seq enables precise characterization of single cell identities in human pre-implantation embryos. (**a**) A workflow showing the generation of scCAT-seq profiles of human pre-implantation embryos. (**b**) Heatmap showing exposure scores of all cells to each signature identified by the NMF clustering of regulatory relationship binary matrix of human embryos. Example genes are shown. **(c)** Regulatory relationships for the indicated genes in single cells of the morula and blastocyst stage. (**d**) Heatmaps showing accessibility deviation (left) and expression level (right) of the indicated TFs. The TFs coloured in green were the ones showing consistent patterns in accessibility and gene expression. (**e**) Immunofluorescence imaging of human morula‐ and blastocyst-stage embryos using the indicated antibodies (left to right: NANOG, SOX17 and merged DAPI/NANOG/SOX17). (**f**) Top and middle panel: Heatmaps showing the accessibility deviation (top) and expression level (middle) of the indicated TFs in single cells of blastocyst-stage embryos. Bottom panel: heatmap showing the expression level of the indicated genes. The TFs coloured in green were the ones showing consistent patterns in accessibility and gene expression.

The transition between cell fates largely depends on TFs, which bind to CREs and recruit chromatin modifiers to reconfigure chromatin structure^15^. Single-cell chromatin accessibility data provides a great opportunity to find the key TFs in individual cells^10, 17^. However, TFs of the same family often share similar motifs, which makes it difficult to determine the key TFs of functional specificity. Previous efforts have proposed computational algorithms to integrate CA and GE data, but the accuracy remains uncertain because the analyses are based on separate multi-omics datasets^16, 17^.

We reasoned that functionally relevant master TFs in each cell type should be determined by integrated omics data obtained by scCAT-seq. We applied chromVAR^29^, a method for inferring TF accessibility with single cell CA data, to compute the deviations of known TFs across all single cells. This method identified TF motifs with high variances (Supplementary Fig. 4d), dividing all single cells into two main groups (Supplementary Fig. 4e), in agreement with the clustering results on regulatory relationships (Fig. 3b). We observed that motifs from the POU-Homebox, SOX-HMG and KLF-zf families showed high deviation score in cells of the group 1, while motifs from GATA-zf and GRHL-CP2 families showed a high deviation score in cells of the group 2 (Fig. 3d). To determine the master TF from each family, we next integrated the expression level of these TFs. Interestingly, we found that the well-known pluripotency factors (such as NANOG, POU5F1, SOX2, KLF4, TBX4), as well as early markers (such as KLF17), both showed relatively high levels of CA and GE in cells of the group 1, whereas other TFs of the same families (such as POU3F1, SOX5, KLF7 and TBX1) showed opposite trends (Fig. 3d). These results are highly consistent with the features of the pluripotent morula cells, which are the main component of group 1. We also found GATA3, but not GATA4 and GATA6, to show a specific role in the group 2, which contains cells from the blastocyst stage. This is in agreement with the important role of GATA3 during differentiation of trophoblast^30^. In addition, we also observed similar results from other TFs of the same families, such as SOX9, HOXD4, MEF2C and GRHL1, suggesting they likely playing critical roles in these two groups (Fig. 3d). Overall, these results suggest that our integrated method could increase the power of discovery of functionally relevant TFs at single-cell resolution.

The blastocyst stage consists of inner cell mass (ICM) and TE lineages. During the maturation of blastocysts, the ICM segregates into pluripotent epiblast (EPI) and primitive endoderm (PE) cells^31^. The number and size of ICM cells vary across blastocysts, and is important for the grading of embryos that determine the success of implantation^32^. Notably, the clustering of both regulatory relationships and TF accessibility deviation showed that 3 (#504, #539, #522) out of the 43 blastocyst cells are similar to morula cells (Fig. 3b). This reveals the pluripotency feature of these 3 single cells in the blastocyst stage and suggests that they might be from ICM cells (hereafter termed ICM-like cells). This result is also supported by our data based on immunostaining in a human blastocyst embryo, which showed a comparable small proportion using the known, lineage-specific markers NANOG (EPI), SOX17 (PE) (Fig. 3e).

We next sought to validate the ICM-like cells by molecular features based on their two omics signatures. It is known that OCT4 is initially expressed in all cells within the ICM, and becomes restricted to the EPI in the late blastocyst^31^. Interestingly, although OCT4 was not a general marker of the blastocyst stage (Fig. 3d), it has a higher deviation score in the 3 single cells compared to other cells in the blastocyst (Supplementary Fig. 4f). Notably, 2 of them (#504 and #539) showed even higher deviations from the other single cell (#522) (Supplementary Fig. 4f), which may describe the segregation into EPI (#504 and #539) and PE (#522) lineages (hereafter termed “EPI-like” and “PE-like” cells).

We next attempted to support this hypothesis by identifying the key TFs in the EPI‐ or PE-like cells. Encouragingly, in addition to enrichment of OCT4, we also observed specific enrichment of the well-known EPI specific regulators, such as NANOG, and KLF17, in EPI-like cells (Fig. 3f), while the PE-like cell showed high activity of the well-known PE regulators, such as SOX17, HNF1B and FOXA2 (Fig. 3f). The other members of the same families (such as SOX9, FOXA1 and HNF1A) are not likely to be the key regulators because of the inconsistent patterns of CA and GE. Further supporting this conclusion, the well-known non-TF markers were also found to be highly specific to each cell type, including GDF3, TGDF1, DPPA2, DPPA5, ARGFX in EPI-like cells and BMP2, PDGFRA, FN1, COL4A1 and LINC00261 in PE-like cells^33^ (Fig. 3f). Although the EPI‐ and PE-like cells are similar to morula cells, the above markers tend to be transcriptionally active in EPI‐ or PE-like cells based on CA and GE profiles. (Supplementary Fig. 4g,h), suggesting distinct pluripotent states in the morula and blastocyst stages. Taken together, these results indicate that our integrated approach can faithfully identify the two distinct subtypes from the same origin. The robustness of scCAT-seq in the precise definition of single-cell identities would be particularly useful for characterization of cells that are rare within complex cell populations.

In summary, our work demonstrates that scCAT-seq is able to provide high resolution epigenomic and transcriptomic portraits of individual cells. We showed that the accessibility levels of both regulatory elements and particular TFs are positively correlated with the GE program. This provides a highly relevant insight into regulatory relationships, one which is not possible based on individual omics profiles. We proposed a method to establish regulatory relationships by linking CREs to the putative target genes, resulting in a larger numbers of high-confidence regulatory interactions compared to state-of-the-art methods. The cell-specific regulatory relationship is a new feature that enables the direct discovery of gene centered 3D regulatory patterns in certain cell populations, thus providing the basis for a more comprehensive study of regulatory mechanisms at the single cell level. Moreover, we generated the first integrated single cell epigenomic and transcriptomic maps during preimplantation embryo development. The robustness of scCAT-seq in the characterization of distinct cell states reveals the great potential of scCAT-seq in faithful identification of new cell types in complex cell populations, which enables a better understanding of developmental abnormalities caused by either genomic variants or environmental influences. Overall, we show that scCAT-seq is a highly promising tool for the joint study of multimodal data of single cells, paving the way to a thorough assessment of regulatory heterogeneity in a variety of clinical applications including pre-implantation screening.

## ACKNOWLEDGEMENTS

We thank all members of the Stem Cell and Development Lab (BGI) for their support and Scott Edmunds, Christian Conrad, Kun Ma and Ying Shan for helpful discussion. We also thank Jijun Cheng, Yuan Long and Feifei Zhang from Shanghai LIDE Biotech Co., Ltd. for technical support. This work was supported by the Strategic Priority Research Program of the Chinese Academy of Sciences (XDA16010100), the Shenzhen Municipal Government of China Peacock Plan (KQTD20150330171505310) and the Shenzhen Engineering Laboratory for Innovative Molecular Diagnostics (DRC-SZ (2016) 884). Longqi Liu is funded by the China Postdoctoral Science Foundation (2017M610553).

## COMPETING FINANCIAL INTERESTS

The authors declare no competing financial interests.

---

**Supplementary Figure 1.**
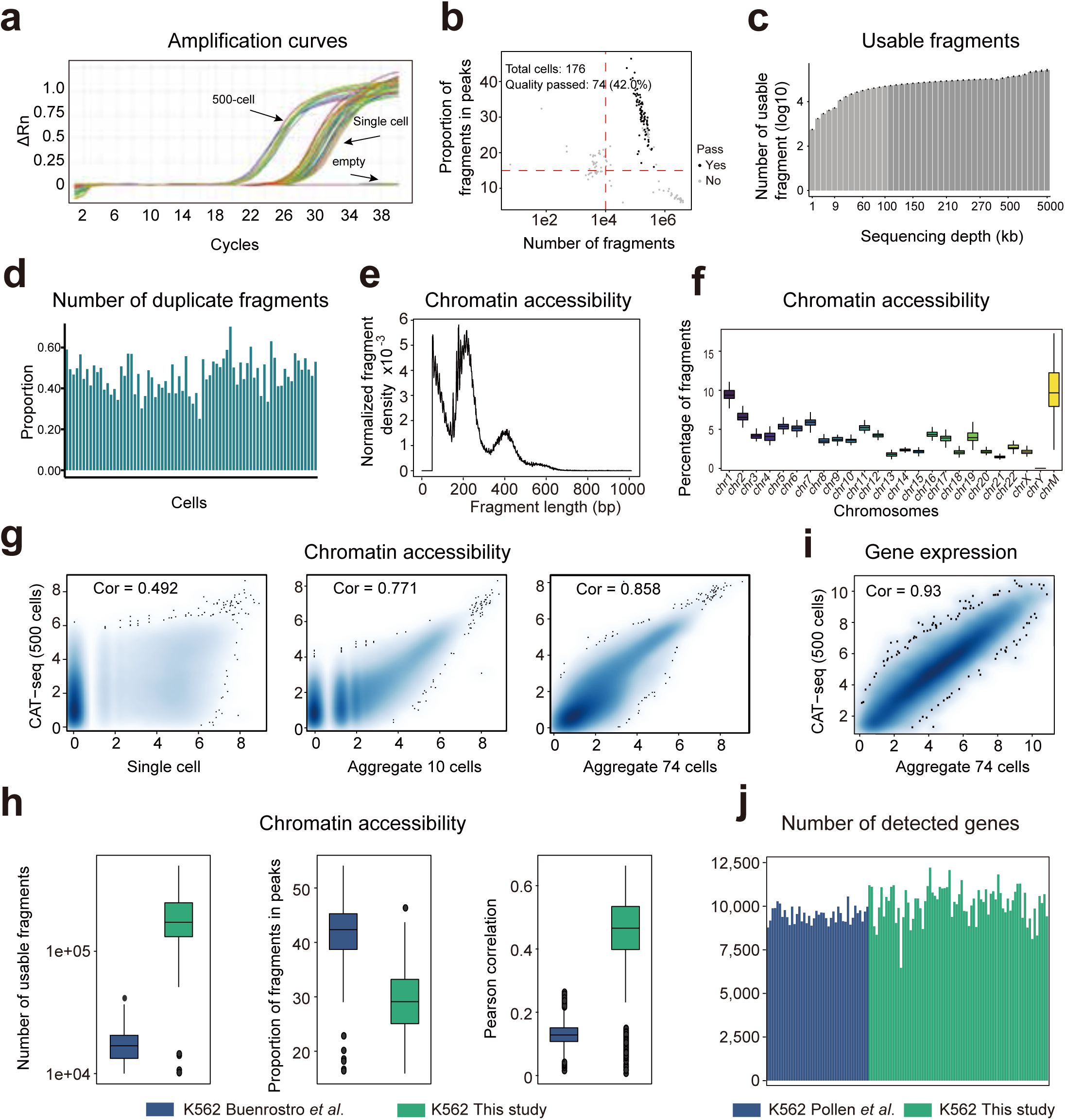
Quality metrics of scCAT-seq data. (**a**) qPCR amplification curve using materials in the bottom of wells after the separation step of the scCAT-seq protocol. Wells containing 0, 1 and 500 cells were analyzed. After the separation step the materials were amplified for 8 cycles using primers targeting the Tn5 adaptor. The PCR product was then purified and amplified by qPCR using primers targeting an accessible region in the human genome. (**b**) K562 scCAT-seq profiles were quality-filtered according to the number of fragments, proportion of fragments within accessible regions and detected gene numbers. **(c)** Bar plot showing the number of usable fragment at the indicated sequencing depths **(d)** Proportion of the duplicate fragments of all K562 single cells at the sequencing depth of 400 kb. (**e**) Size distribution of chromatin accessibility fragments from an example of K562 single cell. (**f**) Percentage of the single cell chromatin accessibility fragments mapped to each nuclear chromosome and the mitochondrial genome. **(g)** Correlation of chromatin accessibility between aggregate chromatin accessibility profiles and CAT-seq profile of 500 cells. **(h)** Comparison of number of usable chromatin accessibility fragments (left), proportion of fragments within the accessible regions (middle) and Pearson correlation coefficients (right) between scCAT-seq and published scATAC-seq profiles. The peaks indicated in middle panel are called based on aggregate profiles. **(i)** Correlation between aggregate gene expression profiles of all single cells and gene expression profiles generated from 500 cells. (**j**) Comparison of the number of detected genes between scCAT-seq and published scRNA-seq profiles.

**Supplementary Figure 2.**
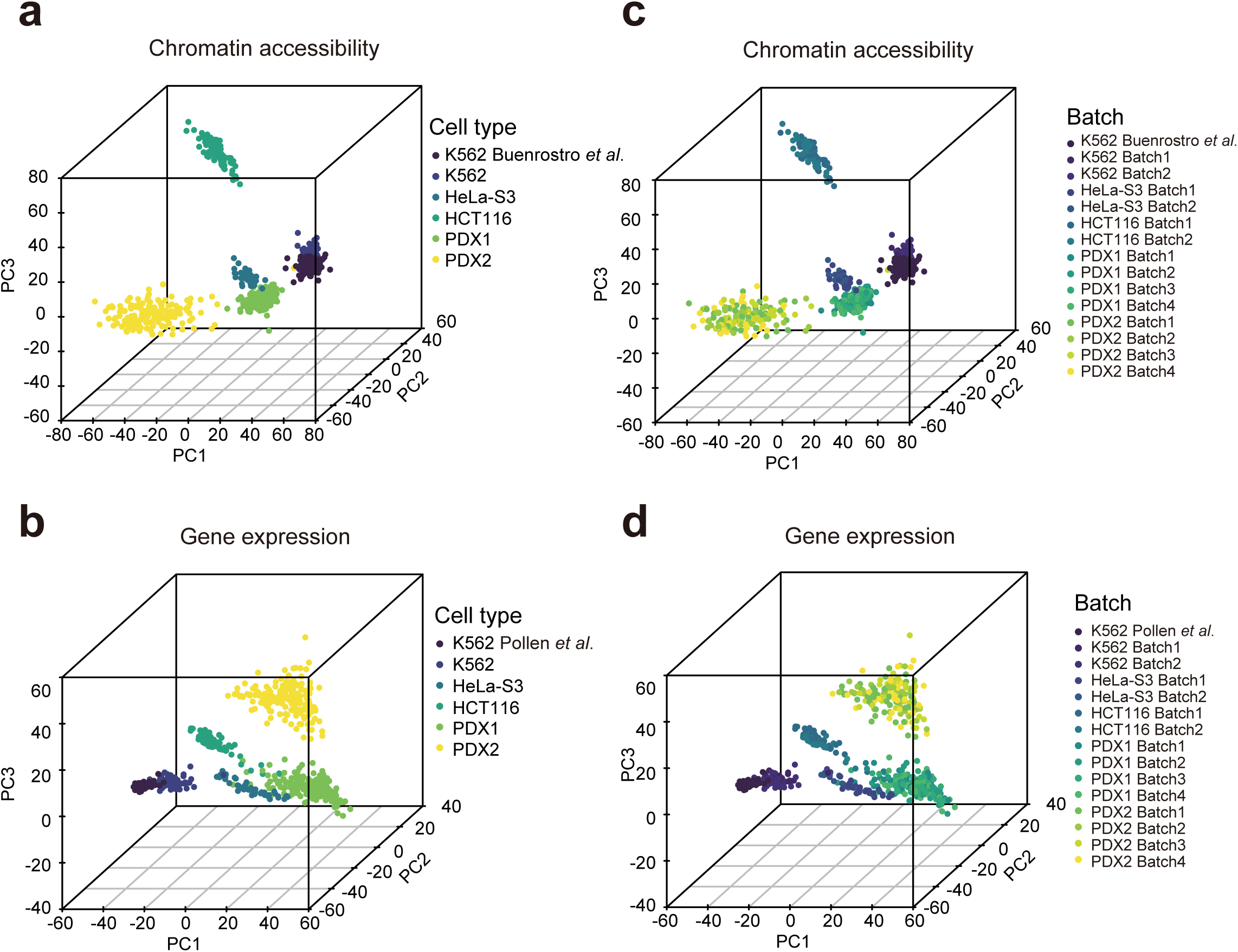
Principle components analysis across diverse techniques and different batches of scCAT-seq profiles. **(a** and **c)** Principle components analysis of different batches of scCAT-seq-generated chromatin accessibility data and published datasets. **(b** and **d)** Principle components analysis of different batches of scCAT-seq-generated gene expression data and published datasets.

**Supplementary Figure 3.**
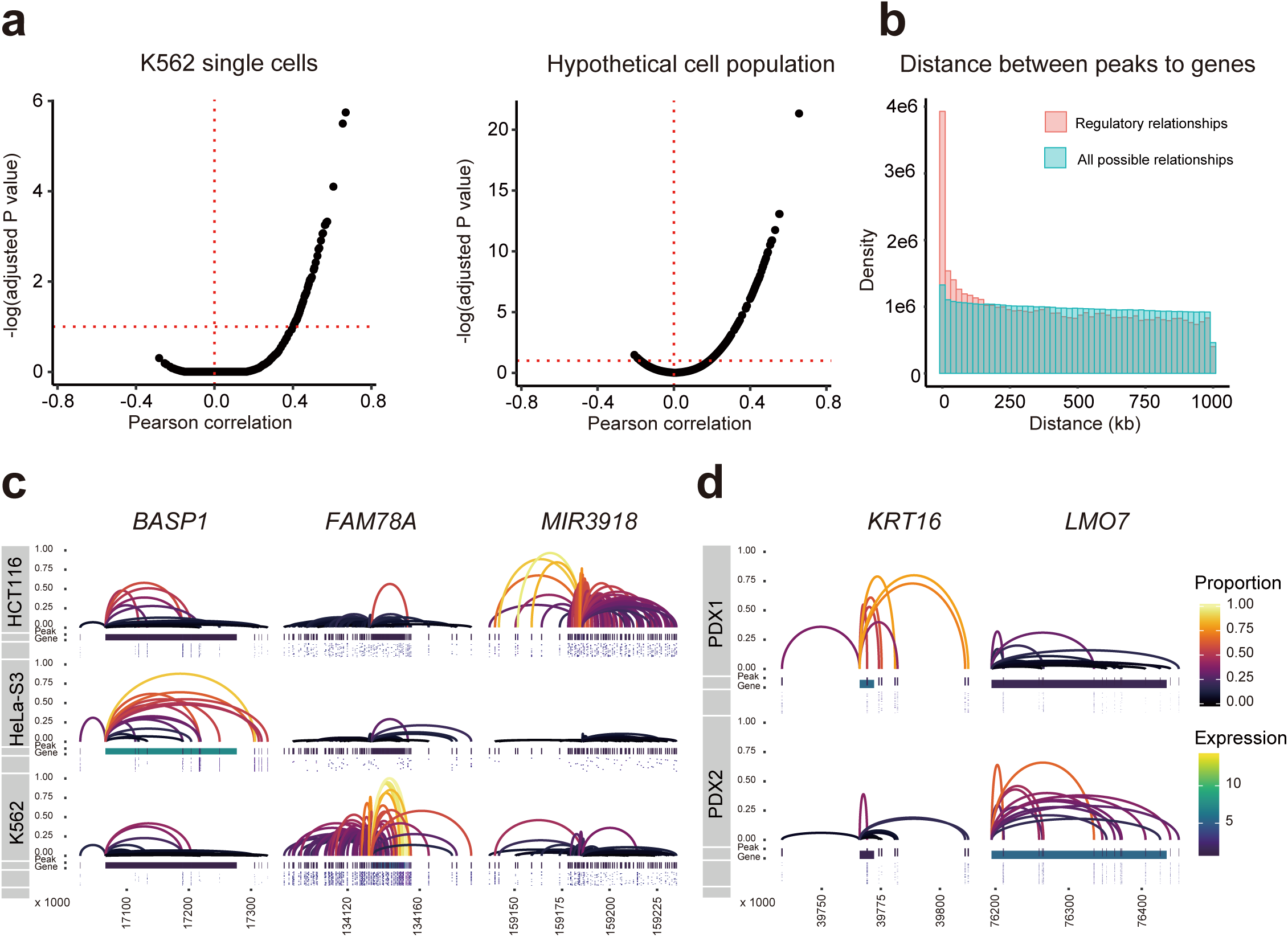
scCAT-seq uncovers the regulatory relationships between CREs and genes. **(a)** Correlation analysis between chromatin accessibility of individual element and the putative gene expression in K562 single cells and hypothetical cell population from the three cell lines. Shown are Pearson correlation coefficients versus the Benjamini-Hochberg adjusted p-value. Significant relationships (adjusted p-value <= 0.05) are above the red dotted line. **(b)** Bar plot showing the density distribution of distances between CREs and genes in regulatory relationships (red) and random relationships (blue) **(c-d)** Regulatory relationships for the indicated genes in single cells of the three cell types **(c)** and two PDX tissues **(d)**.

**Supplementary Figure 4.**
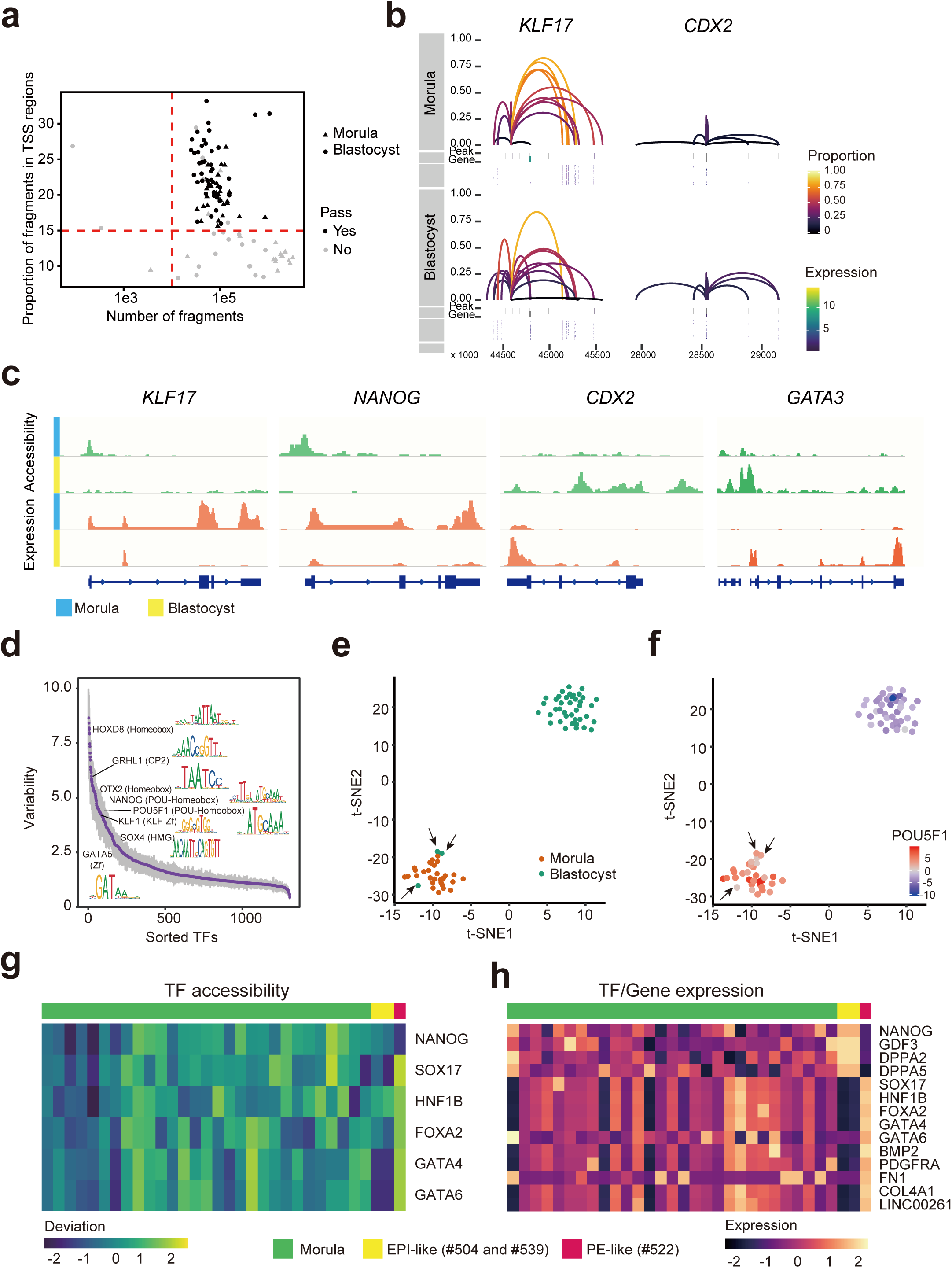
Integrated profiling of chromatin accessibility and gene expression in human pre-implantation embryos. **(a)** Morula and blastocyst scCAT-seq profiles were quality-filtered according to the number of fragments, proportion of fragments within promoter regions and detected gene number. **(b)** Regulatory relationships for the indicated genes in single cells of morula and blastocyst stage. **(c)** Genome browser views of chromatin accessibility and gene expression surrounding the indicated genes. **(d)** Observed cell-to-cell variability of TFs. TF families and motifs are indicated. **(e)** t-SNE plot of TF motif accessibility deviation, colored by the stage of all single cells. **(f)** t-SNE plot colored by accessibility deviation z-score of POU5F1 motif. The three blastocyst cells that are closed to the morula cells are highlighted with the black arrows. **(g)** Heatmaps showing accessibility deviation (left) and expression level (right) of the indicated TFs.

